# pWCP is a widely distributed and highly conserved *Wolbachia* plasmid in *Culex* mosquitoes worldwide

**DOI:** 10.1101/2022.09.14.507914

**Authors:** Amani Ghousein, Jordan Tutagata, Manuel Etienne, Victor Chaumeau, Sebastien Boyer, Nonito Pages, David Roiz, A. Murat Eren, Guillaume Cambray, Julie Reveillaud

## Abstract

Mosquitoes represent the most important pathogen vectors and are responsible for the spread of a wide variety of poorly treatable diseases. *Wolbachia* are obligate intracellular bacteria that are widely distributed among arthropods and collectively represents one of the most promising solutions for vector control. In particular, *Wolbachia* has been shown to limit the transmission of pathogens, and to dramatically affect the reproductive behavior of their host through its phage WO. While much research has focused on deciphering and exploring the biocontrol applications of these WO-related phenotypes, the extent and potential impact of the *Wolbachia* mobilome remain poorly appreciated. Notably, several *Wolbachia* plasmids, carrying WO-like genes and Insertion Sequences (IS), thus possibly interrelated to other genetic units of the endosymbiont, have been recently discovered. Here we investigated the diversity and biogeography of the first described plasmid of *Wolbachia* in *Culex pipiens* (pWCP) in several islands and continental countries around the world—including Cambodia, Guadeloupe, Martinique, Thailand, and Mexico—together with mosquito strains from colonies that evolved for 2 to 30 years in the laboratory. Together with earlier observation, our results show that pWCP is omnipresent and strikingly conserved among *Wolbachia* populations within mosquitoes from distant geographies and environmental conditions. These data suggest a critical role for the plasmid in *Wolbachia* ecology and evolution, and the potential of a great tool for the further genetic dissection or potential manipulation of the endosymbiont.

## Introduction

The widespread intracellular bacterium *Wolbachia* has been at the heart of mosquito biocontrol programs for decades and is now more than ever triggering a surge of interest due to recent discoveries broadly related to its mobile genetic elements (its mobilome). Remarkably, *Wolbachia* is capable of manipulating the reproduction of its host, thereby favoring its own—almost exclusively maternal—spreading. It has also been shown to provide strong protection against the transmission of viral pathogens by mosquitoes (Ant et al., 2018; Nazni et al., 2019; O’Neill et al., 2018). Together, these properties champion *Wolbachia* as one of the promising strategies for vector control worldwide.

Recently, the most common effect of host reproduction manipulation, Cytoplasmic Incompatibility (CI) was found to be associated with the *cifs* genes harbored by *Wolbachia* bacteriophage WO (LePage et al., 2017; Shropshire et al., 2018; Bonneau, Landmann, et al., 2018; Bonneau, Atyame, et al., 2018; Sicard et al., 2021). These genes are part of a so-called Eukaryotic Associated Module (EAM) that presumably aids phage particles to cope with both prokaryotic and eukaryotic cell membranes, as well as cytoplasmic and extracellular host environments (Bordenstein and Bordenstein, 2022, 2016). WO commonly appears as a temperate prophage integrated in the chromosome and while lytic events remains rarely observed (Bordenstein and Bordenstein, 2016; Fujii et al., 2004), it provides a central source of evolutionary innovation and adaptation for the restricted lifestyle of the obligate intracellular endosymbiont (Bordenstein and Bordenstein, 2022; Bordenstein and Wernegreen, 2004; Chafee et al., 2010).

Being an obligate intracellular symbiont, focus on the mobilome of *Wolbachia* spp. has long been restricted to WO, until the discovery of the first *Wolbachia* plasmid—named pWCP for “plasmid of *Wolbachia* in *Culex pipiens”*—opened new perspectives (Reveillaud and Bordenstein et al., 2019). pWCP was originally reported from *Culex pipiens pipiens* specimens from the Mediterranean basin, including Southern France and Northern Africa (Tunisia, Algeria and Turkey). However, the actual distribution and potential variability of pWCP in *Culex* mosquitoes remain unknown. Most pWCP-born genes were initially found exclusively in *Wolbachia* from *Culex quinquefasciatus* (*w*Pip, (Klasson et al., 2008) and/or to the phylogenetic supergroup B-*Wolbachia*. Very recently, several homologous genes were identified in other *Wolbachia* supergroups from *Aedes albopictus* mosquitoes and other insect species, as part of other novel plasmids (Martinez et al., 2022). Similar to pWCP, plasmids of *Wolbachia* endosymbiont *w*AlbA 1 and 2 (namely pWALBA1 and pWALBA2) include a *parA*-like partitioning gene and a *RelBE* toxin–antitoxin system, supporting the likely functional importance of these elements in a plasmid context. In addition to putative phage-like proteins in *w*AlbA1, the authors reported for the first time *cif* genes homologs in the reconstructed plasmid from two reanalyzed *Wolbachia* genomes (Insecta_WOLB1166 and *D. virgifera virgifera*) together with plasmid-like islands located next to WO prophage regions in *O. gibbosus spiders*. These novel data substantiate the idea that interactions between *Wolbachia* mobile genetic elements could enhance the adaptation and innovation capabilities of these endosymbionts. This remains to be further investigated in different mosquito species.

Notably, the presence of pWCP in *Culex* species requires critical attention. Indeed, the *Culex pipiens* complex represents the most widespread mosquitoes around the world (Bhattacharya and Basu, 2016; Aardema, Olatunji, and Fonseca 2022). It is comprised of the tropical species *Culex quinquefasciatus* and the temperate species *Culex pipiens*, itself divided into two subspecies *Cx. pipiens molestus* and *Cx. pipiens pipiens*. Concomitant to its wide distribution, *Culex* species are vectors of numerous pathogens, causing a variety of known diseases that include West Nile Virus (WNV), one of the most commonly transmitted mosquito disease in the United States (Curren et al., 2018), St. Louis Encephalitis Virus (SLEV), Japanese Encephalitis Virus (JEV), Rift Valley Fever (RVF, Gregor et al., 2021) and the emerging virus Usutu (Cook et al., 2018). Furthermore, the combination of ubiquitous distribution, vector competence and opportunistic feeding behavior provides *Culex* mosquitoes with a high capacity to transmit infectious diseases between animal and humans (*i*.*e*., zoonoses), which represents an important threat to human health (Weissenböck et al., 2010). In fact, a large percentage of all newly identified infectious diseases are zoonoses, some of which have the potential to cause global pandemics, as recently demonstrated by the novel coronavirus that causes COVID-19 (Holmes, 2022; Pekar et al., 2022).

Here, we collected *Culex* spp mosquito specimens from several continents and islands around the world including Thailand, Cambodia, Martinique, Guadeloupe, and Mexico together along with laboratory colonies that have evolved for 2 to 30 years in the laboratory (*Culex pipiens molestus* and *Culex quinquefasciatus* SLAB, respectively), and screened for the presence and variability of pWCP in the germline and somatic tissues of these widespread samples.

## Results

### Screening of pWCP in Wolbachia-infected Culex samples

We collected and dissected the ovaries and midguts of field *Culex quinquefasciatus* mosquito specimens from Cambodia, Thailand, Guadeloupe, Martinique, and Mexico (Figure 1a, Supplementary Table 1). In addition, we sampled *Culex pipiens molestus* specimens originating from a colony that we collected in Montpellier (South of France) and maintained in the laboratory for 2 years (2020-2022) as well as a *Culex quinquefasciatus* SLAB samples, which have been kept more than 30 years in lab environment. Our sampling effort including 35 *Culex* specimens aimed to search for pWCP in mosquitoes from both continental and islands areas across the globe, as well as distinct environmental and laboratory settings.

**Figure 1.**
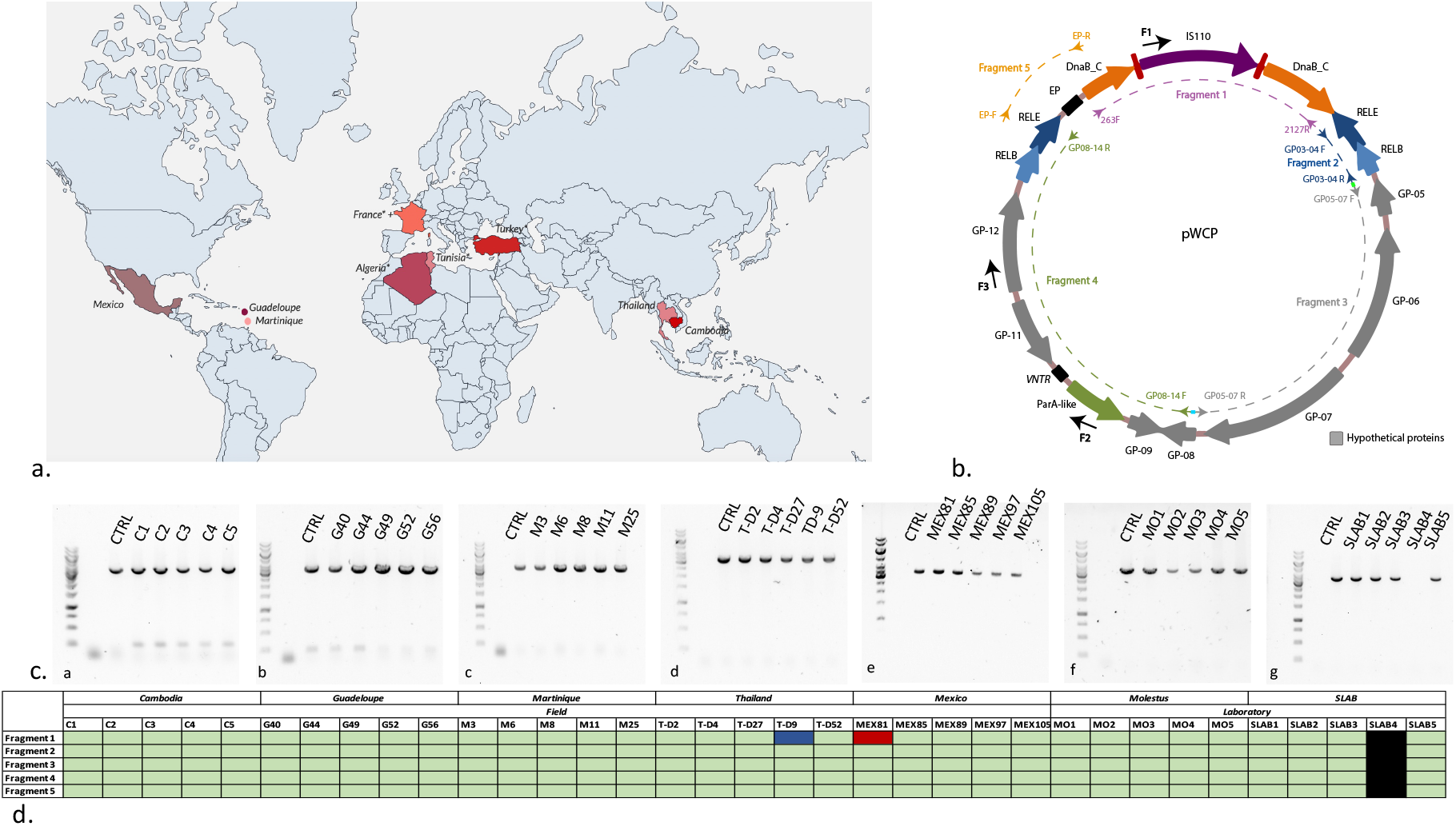
**a. Geographic map with locations of *Culex* sample collection for pWCP screening.** Colors indicate field samples. A “+” indicates laboratory specimen and “ * “ indicates previous observations from (Reveillaud and Bordenstein et al., 2019). **b. Map of pWCP plasmid** (adapted from Reveillaud and Bordenstein et al., 2019). Genes are shown as filled arrows. Couples of PCR primers spanning different regions of the plasmid designed to cover pWCP are shown as smaller arrows. Additional sequencing primers are shown in black arrows on the outer layer. **c. PCR amplification of largest Fragment 4 in studied samples**. A ca. 3451 bp PCR product corresponding to the amplification of Fragment 4 including seven genes of pWCP (GP08, GP09, ParA-like, VNTR, GP11, GP12 and RelBE-2) in most ovary samples collected from different regions. a: Cambodia, b: Guadeloupe, c: Martinique, d: Thailand, e: Mexico, d: Montpellier (molestus) and f: SLAB. **D. Synthetic heatmap of conserved pWCP fragments across samples**. pWCP fragments 1-5 are shown in row and studied samples in column. Green color indicates fragment of the right size, blue if longer, red if shorter. Black corresponds to *Wolbachia*-free sample.

We first confirmed the presence of *Wolbachia* in the ovaries of the different *Culex samples*. A PCR-amplification using specific primers targeting the 16S ribosomal RNA gene of *Wolbachia* clearly showed that all collected *Culex* specimens were infected by *Wolbachia* (Supplementary Figure 1, Supplementary Table 2), except for one of the five SLAB samples. Although a sensitivity issue due to very low *Wolbachia* density could possibly explain the lack of PCR amplification for this sample, it is also possible *Wolbachia* was not transmitted in SLAB4 in this laboratory colony.

We then screened for the presence of pWCP in the 34 *Wolbachia*-infected *Culex* samples. For that purpose, we designed and used five sets of primers spanning regions of the plasmid to eventually cover the entirety of pWCP (Figure 1b). Our first PCR amplification (aiming to amplify to the *DnaB_C* gene and the IS110 region, hereafter Fragment 1, successfully produced the expected PCR product for most samples (1800 bp; 33/34 samples) except for TD-9 and MEX81 showing a larger (ca. 3000 bp) and a shorter (ca. 400 bp) DNA fragments, respectively (Supplementary Figure 2).

A second amplification with primers designed to amplify GP03-04 genes of pWCP produced of band of the expected size of ca. 609 bp for each *Wolbachia*-infected *Culex* samples (Fragment 2, Supplementary Figure 3). For a third pWCP amplification, we designed primers to amplify the region covering the three hypothetical proteins GP05, GP06 and GP07 of pWCP (Fragment 3, Supplementary Figure 4). We detected a ca. 2500 bp fragment which confirmed the presence of the three genes in all *Wolbachia*-infected samples. Our fourth PCR amplification with primers designed to amplify seven genes of the pWCP (GP08, GP09, ParA-like, VNTR, GP11, GP12 and RelBE-2) generated a PCR product of 3451 bp for all the above samples (Fragment 4, Figure 1c). Finally, we designed and used primers to amplify the *RelEB*, EP and *DnaB_C* region which produced a band of 678 bp in a fifth amplification (Fragment 5, Supplementary Figure 5).

In the same manner, we screened for the presence of pWCP in the midgut samples of the same *Wolbachia*-infected individuals using a selection of primers in order to investigate whether the plasmid also occurred in mosquito somatic tissues. We only observed a faint band in 3 out of 14 representative samples (that is, two per geographic locations) for all fragments which is likely due to a relatively low abundance of the plasmid in non-germinal organs (Supplementary Figure 6).

Plasmid is therefore present in all sample with only minor topological variation, 2 out of 35 have different sizes. Successful PCR and similar size suggest high gene content conservation.

### *Estimating pWCP variability in Culex* spp *across the globe*

We next sought to quantify the extent of sequence diversity present in the pWCP genome across different geographical regions and conditions. For this, we used Sanger sequencing of the PCR products obtained above in the 34 *Wolbachia*-infected *Culex* samples and aligned the resulting sequences against the reference pWCP (see Supplementary Table 3 for sequencing primers). We observed no variation in Fragment 1 (925 nts, Alignment 1A; 918nts, Alignment1B), Fragment 2 (533 nts, Alignment 2), Fragment 3 (712 nt, Alignment 3), Fragment 4 (970 nts, Alignment 4A, except for the variable number tandem repeat (VNTR) region as expected and described in Alignment 4B below, 998 Alignment 4C and 901 Alignment 4D), nor Fragment 5 (613, Alignment 5) where each of the sequences from the set of *Culex* samples were 100% identical, revealing a high level of conservation across continents (Figure 1d).

Sequence of the smaller Fragment 1 of sample MEX81 matched the reconstructed sequence of *DnaB_C* gene of pWCP upon excision of the IS110 transposase with a 3 nucleotides scar (Alignment 1C). Conversely, BLASTp searches, using ORFs identified in the longer Fragment 1 from sample TH-D9, allowed us to identify strong similarity (97% similarities over 118 residues) to an IS630-related transposase from various *Wolbachia* of *Culex quinquefasciatus* Pel strain genomes (*e*.*g*. NZ_DS996942.1), thus hinting at the probable insertion of another IS within the IS110 of pWCP (Alignment 1D).

The variable number tandem repeat (VNTR) region of pWCP, identified as pp-hC1A_5, and used to genotype different strains of *Wolbachia* (Petridis and Chatzidimitriou, 2011) was shown to contain a number of repeats that varies across individuals (Reveillaud and Bordenstein et al., 2019). We here observed an insertion of 16 bp for each sample except TH-D2 and MEX85 as compared to the original pWCP sequence together with punctual mutations in all samples (Alignment 4B), further revealing the variable nature of the VNTR region at the level of each individual.

Overall, these results hint at an unexpectedly high degree of conservation of pWCP around the globe.

## Discussion

We screened for the presence and variability of pWCP among 30 *Culex quinquesfasciatus* and 5 *Culex pipiens molestus* specimens sampled across the European, North American and Asian continents as well as from several islands. Our collection included freshly collected wild specimens as well as samples originating from mosquito strains maintained in the lab for two to more than 30 years. PCR and Sanger sequencing results indicate that pWCP is widely distributed and highly conserved among *Culex* spp worldwide. We observed identical pWCP fragments in nearly all *Culex* mosquito specimen studied herein except for the VNTR region, as expected for this variable region with high repeat content. The transposase IS100 was found in all samples (except for sample MEX 81) suggesting an ancient, and perhaps functional insertion event in the *DnaB-C* gene borne by pWCP. It could suggest that i) the *DnaB-C* gene is disrupted and not functional, or more probably that ii) both *DnaB-C* gene subunits can re-assemble, as the whole gene does not appear as degenerated amongst samples. Our results indicate that the presence of pWCP is not incidental and that its ordered and highly conserved genes are likely to be functional. Of note, one sample from Thailand (D9) showed a larger *DnaB*_*C* gene and IS fragment. Blast analyses of the region resulted in matches to both IS630 and IS110 of *w*Pip, which are frequently found in WO genomes (Kent and Bordenstein, 2010) and further suggest some potential contact and infection events between the different types of *Wolbachia* mobile genetic elements. Nevertheless, in contrast to the newly identified pWALBA1 and pWALBA2 in the invasive species *Aedes albopictus*, pWCP appears as a small and conserved element of ca. 9500 bp in the widespread common house mosquitoes *Culex pipiens/quinquefasciatus* across the globe.

It is striking that completely identical pWCP were found in both field and laboratory settings. These findings highlight that the plasmid is maintained across generations in very different environments and associated selective pressures. In line with this, we observed the presence of pWCP in the ovaries as well as the midguts of *Culex* specimens, suggesting pWCP follows the trajectories of *Wolbachia* transmission from germlines to somatic tissue, where its role remains unknown. Overall, the presence of a conserved pWCP at large scale, covering four continents including North Africa (Reveillaud and Bordenstein et al., 2019), Europe, America and Asia reinforces the notion that this mobile genetic element plays an important role in *Wolbachia* biology. These data converge with the remarkably high average nucleotide identity across *Wolbachia* genomes reconstructed from Southern France and *w*Pip Pel reference genome originally from Sri Lanka (99.1–99.98%, Reveillaud and Bordenstein et al., 2019) which suggested a high degree of core and essential genome conservation across individuals. In addition, the detection for pWCP in three species of the *Culex pipiens* mosquito complex, that is *Cx. quinquefasciatus* and *Cx. pipiens* (var. *pipiens* and var. *molestus*) that are vectors of pathogens opens important perspectives for novel vector control strategies that may have great impact in the battle against pathogens spread from diverse mosquito species.

## Materials and methods

### Mosquito sampling

Collection and dissection of mosquito specimens was performed as described in (Reveillaud and Bordenstein et al. 2019) following a standardized protocol for each location. Briefly, we collected mosquitoes using a carbon dioxide mosquito trap (BG-Sentinel with BG lure or CDC trap baited with carbon dioxide) or an aspiration device and transported them alive to the laboratory directly afterward. Females were identified at species-level, anesthetized by incubation at -20°C for 4 minutes (min), surface-sterilized with ethanol 96% for 1 min, quickly rinsed with sterile PBS to avoid DNA fixation by ethanol, transferred in a drop of sterile PBS deposited on a sterile microscope slide and dissected using sterilized tweezers. Ovaries and midgut from each single mosquito were separated and stored in sterile buffer to preserve them until further processing.

### DNA extraction and PCR amplification

DNA from each organ was extracted using a Qiagen DNeasy blood and tissue kit according to manufacturer’s instructions after rinsing samples with 1000 ul PBS and centrifugation at 12.000 g at 15°C for 15min. Dna was quantified using Qubit (Supplementary Table 1). We used primer CQ11F2 (5’-GATCCTAGCAAGCGAGAAC-3’) and pipCQ11R (5’-CATGTTGAGCTTCGGTGAA-3’) and molCQ11R (5’-CCCTCCAGTAAGGTATCAAC-3’) to confirm the taxonomy of *Culex pipiens pipiens* versus *molestus* (Fonseca and Bahnck, 2006). The presence of *Wolbachia* was monitored by amplifying the 16S rRNA gene using Wspec F and R primers, while the presence of pWCP was investigated by using a set of five specific primers as follows: 263F and 2127R for fragment 1, GP-03-04F and GP-03-04R for fragment 2, GP05-07F and GP05-07R for fragment 3, GP08-14F and GP08-14R for fragment 4 and EP-F and EP-R for fragment 5. All primer sequences are listed in Supplementary Table 2.

PCR amplifications were performed using 1 ng of DNA as template material, 5 μL of 5x reaction buffer, 1 μL dNTPs (10 mM), 0.25 μL of Phusion DNA polymerase (NEB) and nucleic-acid-free water to a final volume of 25 μL. PCR products were electrophoresed on a 0.8% agarose gel to determine the presence of the desired size product of the amplified DNA, using GeneRuler 1 kb (thermoscientific SM0313) as a ladder. DNA from an ovary *Culex pipiens pipiens* sample collected in Montpellier, France (hereafter Cx1) was used as positive control for PCRs and gels. A negative control devoid of template material was also run within each set of reaction.

### Sanger sequencing

We Sanger-sequenced all PCR amplicons obtained from two randomly chosen samples from each geographic location and show one representative sample for each after checking for the lack of intra-site variation. PCR products were purified using Monarch PCR & DNA Cleanup Kit (NEB) for all fragments, except Fragment 1. Since the PCR for fragment 1 showed substantial non specific bands, the band at ca. 1800 bp was excised and processed using Monarch DNA Gel Extraction kit (NEB). Purification kits were used according to manufacturer’s instructions. The purified PCR products were premixed with different corresponding primers and sent for sequencing using ‘name of the sequencing product’ (Eurofins).

## Data availability

Sequencing data for pWCP is available at https://zenodo.org/deposit/7039954#.

## Acknowledgments

We thank Ana Rivero, Arnaud Berthomieu, Olivier Duron and Hans Schrieke for their help getting access and raising laboratory *Culex* colonies, as well as Gilbert Legoff for dissection training to JT. We are also thankful to Maria José Tolsá Garcia, Paola Martínez Duque and Roger Arana; Kimhuor Suor, Moeun Chhum, Kalyan Chhuoy, and Sony Yean; Fabrice Sonor, Marie-Michelle Clemente and Marie-Julie Sellaye; the team of the entomology and Malaria Elimination Task Force departments of the Shoklo Malaria Research Unit for their contribution to sample collection and dissection of samples in Mexico, Cambodia, Martinique and Thailand, respectively. The Shoklo Malaria Research Unit is part of the Mahidol Oxford University Research Unit, supported by the Wellcome Trust of Great Britain and this research was funded in part by the Wellcome Trust [220211]. For the purpose of Open Access, the author has applied a CC BY public copyright license to any Author Accepted Manuscript version arising from this submission. This work was supported by the ERC RosaLind Starting Grant “948135” to JR.

## Authors contributions

AG performed molecular biology work, analyzed the data and wrote the manuscript. JT, VC, NP performed mosquito collection and dissection. ME, SB and DR coordinated mosquito sampling. AME analyzed the data and wrote the manuscript. GC coordinated molecular biology experiments, analyzed the data and wrote the manuscript. JR conceived and coordinated the study, analyzed the data and wrote the manuscript. All authors have read, contributed, and approved the final version of the manuscript.

