## Supplementary Material for "pWCP is a widely distributed and highly conserved *Wolbachia* plasmid in *Culex* mosquitoes worldwide"

**Supplementary Table 1.** Collected samples together with their associated metadata including individual number, dissected organ, storage buffer, location, field vs. lab settings, date of collection, feeding status, gravid status, person who realized dissection, DNA concentration.

**Supplementary Table 2.** Primers and conditions used for the PCR amplification of the five pWCP regions. Colored nucleotides (nt) indicate overlapping nt between fragments.

**Supplementary Table 3.** Primers used for the sequencing of the 5 pWCP regions and the corresponding alignments

**Alignment 1A.** Sanger sequencing of Fragment 1 for one sample per location using primer 2127R.

**Alignment 1B.** Sequence of Fragment 1 for one sample per location using primer F1.

**Alignment 1C**. Sequence of Fragment 1 for sample MEX81 using primer 2127R.

**Alignment 1D.** Sequence of Fragment 1 for TH-D9 using primer F1.

**Alignment 2.** Sanger sequencing of Fragment 2 for one sample per location using primer GP03-04F.

**Alignment 3.** Sanger sequencing of Fragment 3 for one sample per location using primer GP05-GP07R.

**Alignment 4A.** Sanger sequencing of Fragment 4 for one sample per location using primer GP08-14F

**Alignment 4B**. Sanger sequencing of Fragment 4 for one sample per location using primer F2.

**Alignment 4C.** Sanger sequencing of Fragment 4 for one sample per location using primer F3.

**Alignment 4D.** Sanger sequencing of Fragment 4 for one sample per location using primer GP08-14R.

**Alignment 5**. Sanger sequencing of Fragment 5 for one sample per location using primer EP-F.

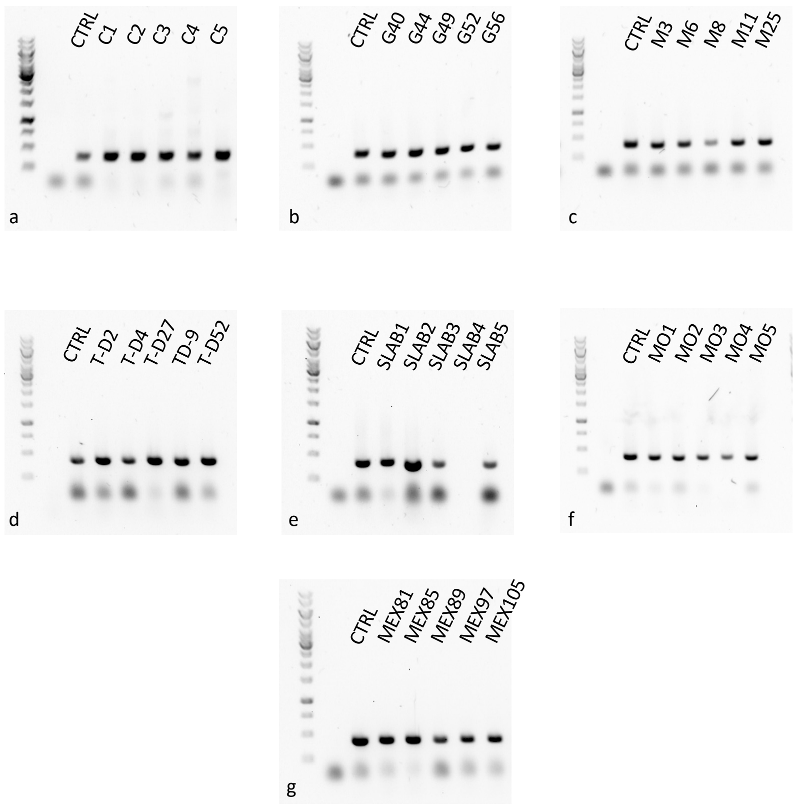


**Supplementary Figure 1**. A ca. 438 bp PCR product corresponding to the amplification of 16S rRNA gene in most ovary samples collected from different regions. a: Cambodia, b: Guadeloupe, c: Martinique, d: Thailand, e: SLAB, f: Montpellier (molestus) and g: Mexico

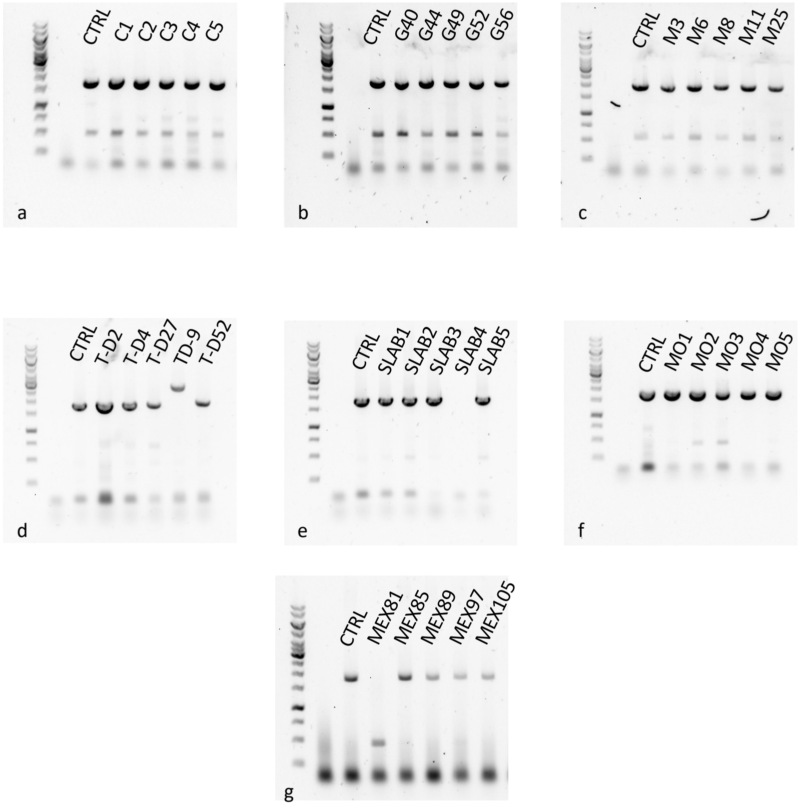


**Supplementary Figure 2.** A 1800 bp PCR product corresponding to the amplification of Fragment 1 of pWCP including *DnaB_C* gene and IS110 in most ovary samples collected from different regions.a: Cambodia, b: Guadeloupe, c: Martinique, d: Thailand, e: Slab, f: Montpellier and g: Mexico. The lower band in several samples of ca. 500 bp corresponds to the amplification of *DNAb_C* gene from *Wolbachia* genome as previously observed in (Reveillaud et al., 2019).

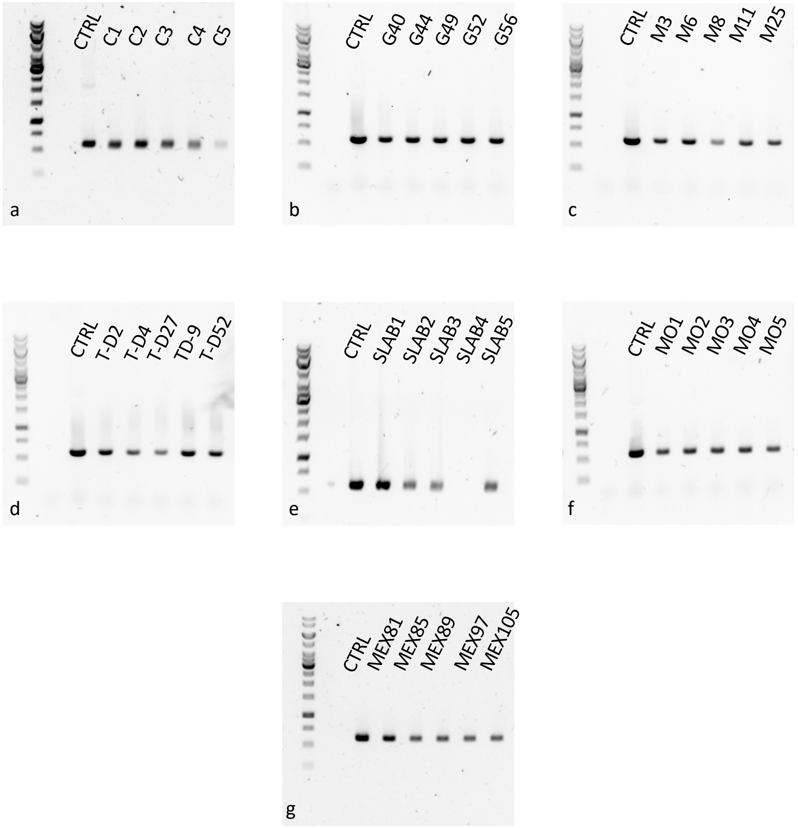


**Supplementary Figure 3**. Amplification of a 609 bp product (Fragment 2) covering genes GP03-GP04 of pWCP in most ovary samples collected from different regions. a: Cambodia, b: Guadeloupe, c: Martinique, d: Thailand, e: SLAB, f: Montpellier (molestus) and g: Mexico.

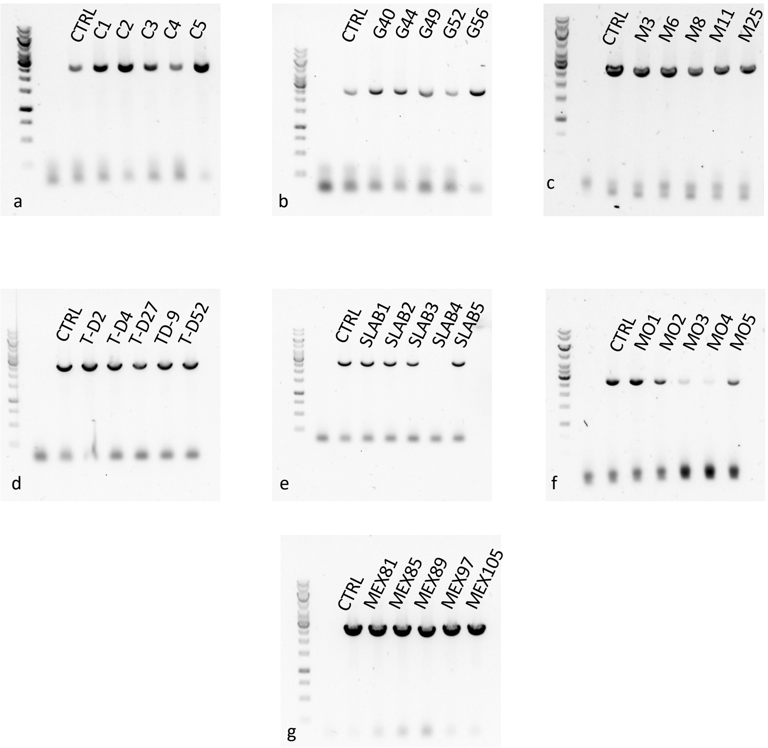


**Supplementary Figure 4**. A ca. 2500 bp PCR product corresponding to the amplification of Fragment 3 including GP05, GP06 and GP07 in most ovary samples collected from different regions. a: Cambodia, b: Guadeloupe, c: Martinique, d: Thailand, e: SLAB, f: Montpellier (molestus) and g: Mexico

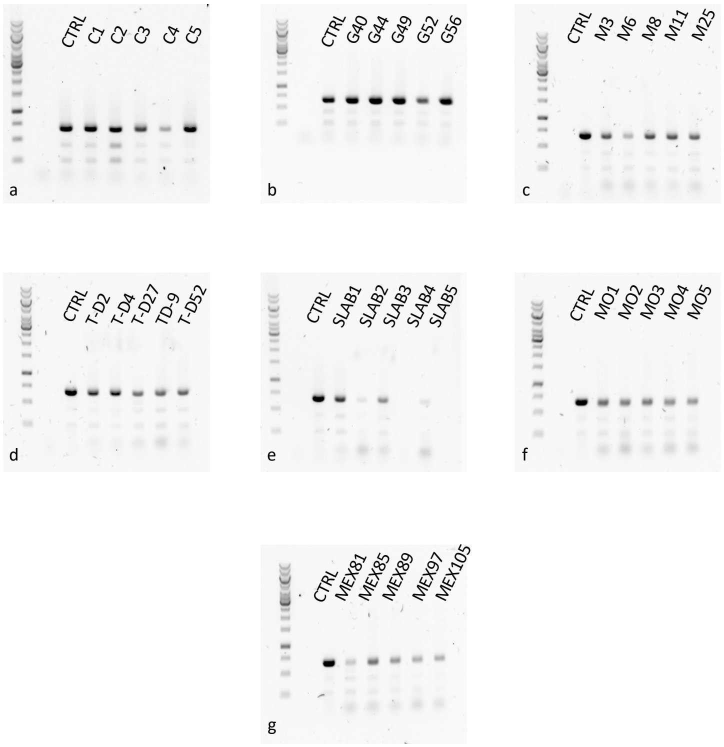


**Supplementary Figure 5**. A ca. 678 bp PCR product corresponding to the amplification of Fragment 5 including the *RelBE*, EP and *DnaB_C* region of pWCP in most ovary samples collected from different regions. a: Cambodia, b: Guadeloupe, c: Martinique, d: Thailand, e: SLAB, f: Montpellier (molestus) and g: Mexico

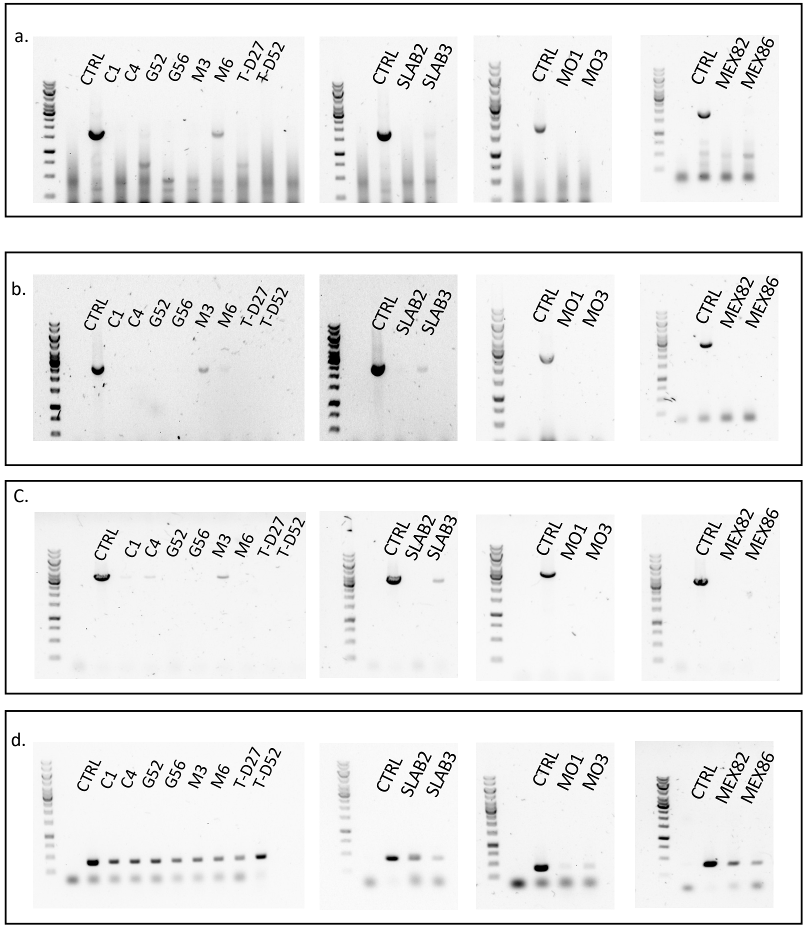


**Suppplementary Figure 6**. Screening of pWCP in midgut samples. **a**. Amplification of a ca. 1800 bp region that corresponds to *DnaB_C* gene with IS (Fragment 1). **b.** Amplification of a ca. 2500 bp ($3^{rd}$ fragment) region that correspond to GP05, GP06 and GP07. **c.** Amplification of a ca. 3451 bp ($4^{th}$ fragment) region that corresponds to seven genes of the pWCP (GP08, GP09, *ParA*-like, VNTR, GP11, GP12 and *Rel*BE-2). **d.** Amplification of the 16S rRNA gene using specific *Wolbachia* primers.
